# Establishment and Thorough External Validation of a FTIR Spectroscopy Classifier for *Salmonella* Serogroup Differentiation

**DOI:** 10.1101/2025.03.18.643663

**Authors:** Helene Oberreuter, Miriam Cordovana, Martin Dyk, Jörg Rau

## Abstract

As one of the most relevant food-borne pathogens, the reliable detection, confirmation and fine-typing of *Salmonella* strains is very important. *Salmonella* serotype determination by rabbit antisera posts the worldwide-accepted standard but is labor intensive, costly and needs extensive experience. As an alternative, successful discrimination between strains of different serogroups by FTIR spectroscopy has been developed before for various bacterial groups. In the current study, firstly a FTIR Classifier operating on an IR Biotyper^®^ spectrometer (Bruker, Germany) was designed to distinguish between *n=*36 different *Salmonella* serogroups. A FTIR classifier is an AI-based tool used in FTIR spectroscopy to analyze and classify different materials based on their infrared spectra.

Secondly, the differentiation performance of this classifier was determined by a thorough external single-lab validation carried out in line with the Guidelines for Validating Species Identifications Using MALDI-ToF-MS issued by the German Federal Office of Consumer Protection for a targeted identification: The most common *Salmonella* serogroups in Europe, serogroups O:4 (B), O:6,7 (C1), O:8 (C2-C3) and O:9 (D1) were chosen as target parameters and validated using a total of *n=*1039 infrared absorbance spectra from a total of *n=*167 strains pertaining to *n=*39 serogroups. In summary, serogroups O:4, O:6,7 and O:9 perfectly met the adapted guideline requirements and resulted in a >99% inclusivity each. Serogroup O:8 arrived at a 96.1% true-positive rate due to one deviating strain. This validated classification method can thus be used in routine analysis for quick and easy differentiation of the most common *Salmonella* serogroups in food surveillance. In addition, using the cluster analysis tools of the IR BT^®^, a preselection of isolates before subjecting them to thorough serotyping decreases the workload in current routine analyses.

## 1 Introduction

*Salmonella* is an important food pathogen and salmonellosis the second most commonly reported gastrointestinal infection in Europe [1]. Nontyphoidal salmonellosis usually manifests itself by a self-limiting diarrhea possibly accompanied by a fever that occurs within 6 to 72 hours after ingestion of the living cells [2]. While salmonelloses occurred much more frequently in the past, the use of live vaccine strains of *Salmonella* (*S.*) Enteritidis and *S.* Typhimurium on breeding and laying hens has helped to greatly decrease the human case frequency [3]. Nevertheless, there are still ∼91000 human illness cases occurring in Europe per year [4] and about 1.3 million annual cases in North America [4]. A continuing frequent carrier of salmonella are reptiles which usually don’t betray their colonization by any illness symptoms [5]. In addition, salmonella bacteria occur widely throughout the environment and are frequently isolated from untreated animal food sources such as meat, eggs and milk in their raw condition. In addition, contaminated fresh produce or spices have also been the sources of outbreaks [2]. Several outbreaks with contaminated ready-to-eat sprouts have been reported where these bacterial germs germinate just as effectively in the warm and moist growth environment as the plant seedlings themselves [6]. Therefore, a quick and simple identification of presumptive colonies is still of high importance.

At the species level, *Salmonella* strains are taxonomically organized into two species, namely *S. enterica* and *S. bongori,* the former of which has been subdivided into six subspecies *S. enterica* ssp. *enterica*, ssp. *salamae*, ssp. *arizonae*, ssp. *diarizonae*, ssp. *houtenae* and ssp. *indica* [7]. While this number is still straightforward, at the serotype level, more than 2600 *Salmonella* serovars are currently recognized [8]. Of these, more than 1500 serovars belong to the subspecies *enterica*, 99% of which may cause infections in animals and humans [9]. The gold standard for *Salmonella* typing is the serotype determination using rabbit antisera. For any given strain, this fine-typing method results in a 3-unit combination code starting with the serogroup indicated by the letter O, followed by two flagellar antigen units H1 and H2. Currently, the 2600 serovars currently recognized have been grouped into 46 O-groups [7].

At the CVUAS strain collection, there are more than *n=*2600 salmonella field strains out of *n=*36 O-groups available deep-frozen which have been isolated from both food products and veterinary samples of domestic, farm and zoo animals. Selected representatives from both the isolate collections at CVUAS and the infrared instruments’ manufacturer Bruker were first used to establish a Fourier Transform infrared spectroscopy (FTIR) classifier that would enable the allocation of unknown salmonella sample isolates to the respective serogroups: the so-called *Salmonella serogroup classifier v3*. Secondly, a different sample set was used to perform an external validation of the FTIR classifier with respect to its capacity for serogroup discrimination.

FTIR spectroscopy is a physico-chemical tool that enables discrimination between different microbial species, subspecies or even serogroups. In principle, dried cells are subjected to infrared light in the mid-infrared region. The light absorption is recorded and the resulting spectra are further treated, e.g. normalized and differentiated to the first or second derivative to enable their further usage. Based on databases consisting of these pre-treated spectra, methods using artificial neural networks can be developed that enable discrimination between the microbial target units. Concerning its handling complexity, FTIR is a comparatively simple and cost-manageable method and its suitability for bacterial typing has been demonstrated before (for reviews, see [10–13]). Specifically, the differentiation between the highly similar *Salmonella* serogroups and serovars has been investigated previously [14–26]. However, the study under consideration is a report on outlining the construction setup of a commercial salmonella IR-Biotyper^©^ (IR BT^©^) classifier and its thorough and formalized external validation carried out in a single-lab.

In this work, firstly, the establishment of the commercially available classifier *Salmonella serogroup classifier v3* is described. The classifier was developed using a support vector machine algorithm included in the IR Biotyper® software and aims to distinguish between *n=*36 salmonella serogroups including the four serogroups containing the serovars *S.* Typhimurium (O:4), *S.* Infantis (O:6,7), *S.* Newport (O:8) and *S.* Enteritidis (O:9) which are most commonly occurring of all the food and veterinary *Salmonella* isolates at the CVUAS lab. Additionally, a further class, named “Others”, is included, comprising both strains of other than the above-mentioned 36 serogroups and strains exhibiting a rough phenotype without any somatic antigens.

Secondly, a structured external single-lab validation of this classifier was carried out according to the adapted national guidelines for MALDI-ToF MS validations [27, 28] as demonstrated previously for the differentiation of *Listeria monocytogenes* serogroups [29]. The *Salmonella* classifier was challenged with a set of *n=*167 different strains belonging to *n=*39 different serogroups. In addition, the *Salmonella serogroup classifier v3* was tried with a group of *n=*16 salmonella strains from lab proficiency tests performed in our lab from 2006 until 2024.

## 2 Materials and Methods

### 2.1 Strains under investigation including species confirmation by MALDI-ToF MS

All isolates used for the validation had been originally isolated at CVUA Stuttgart from food and animal samples according to ISO 6579 [30], followed by subsequent MALDI-ToF MS analysis for formal confirmation on the species level. The MALDI-ToF MS analysis was done on the MALDI Biotyper system, consisting of an LT microflex mass spectrometer and the Compass software, combined with database version K (all Bruker Daltonics, Bremen, Germany) in accordance with the certified workflow for *Salmonella* species confirmation [31]. Prior to their utilization in the current study, each of these isolates’ identification had been confirmed or determined, respectively, by the National Reference Laboratory (*NRL Salmonella*) at the German Federal Institute for Risk Assessment (BfR) together with their respective serotype information.

The list of isolates which were used for the establishment of the commercially available *Salmonella serogroup classifier v3* compiles *n=*158 isolates from *n=*36 different serogroups. Its emphasis lies on serogroups O:4, O:6,7, O:8 and O:9, out of which 75 different strains, i.e. 10-30 each, were included, respectively. 83 strains were arbitrarily chosen pertaining to serogroups O:3 and O:11 through O:65 including rough forms without any positive antisera reaction to comprise a challenging outgroup: the intention was to set up a comparatively large and diverse set of strains in the background to challenge the classifier with real-(lab)life isolates. As strains from two different serotypes were arbitrarily selected per serogroup, this outgroup was referred to as the “Noah’s Ark” strain set.

Likewise, the list of isolates which were used for the external validation of the *Salmonella serogroup classifier v3* compiles *n=*167 isolates from the same *n=*39 O-groups (Tab. 1). None of these had been used by the manufacturer to compile the classifier. With respect to the most common serogroups O:4, O:6,7, O:8 and O:9, *n=*25 different strains were included in the validation set each (see Tab. 1).

**Table 1:**
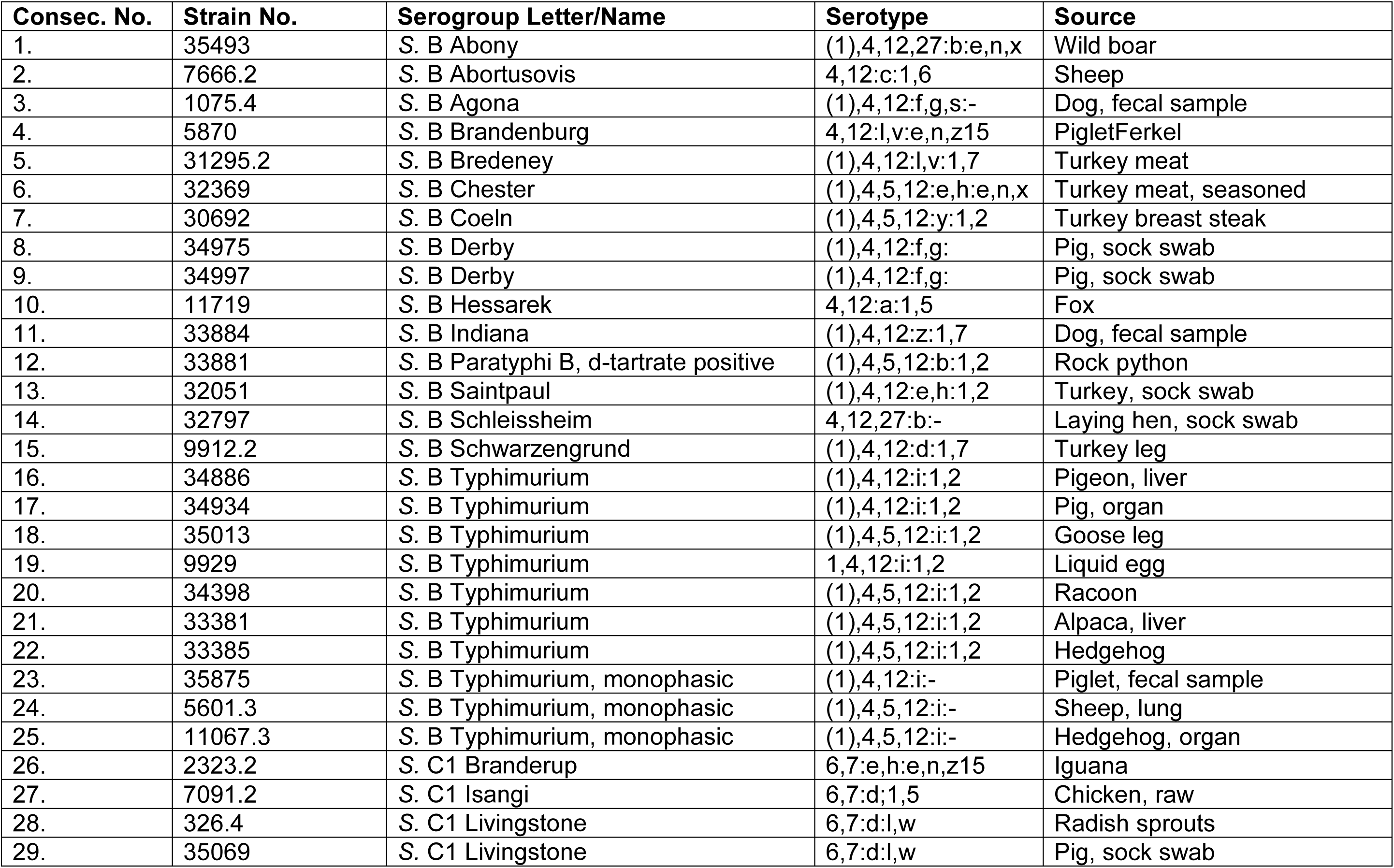
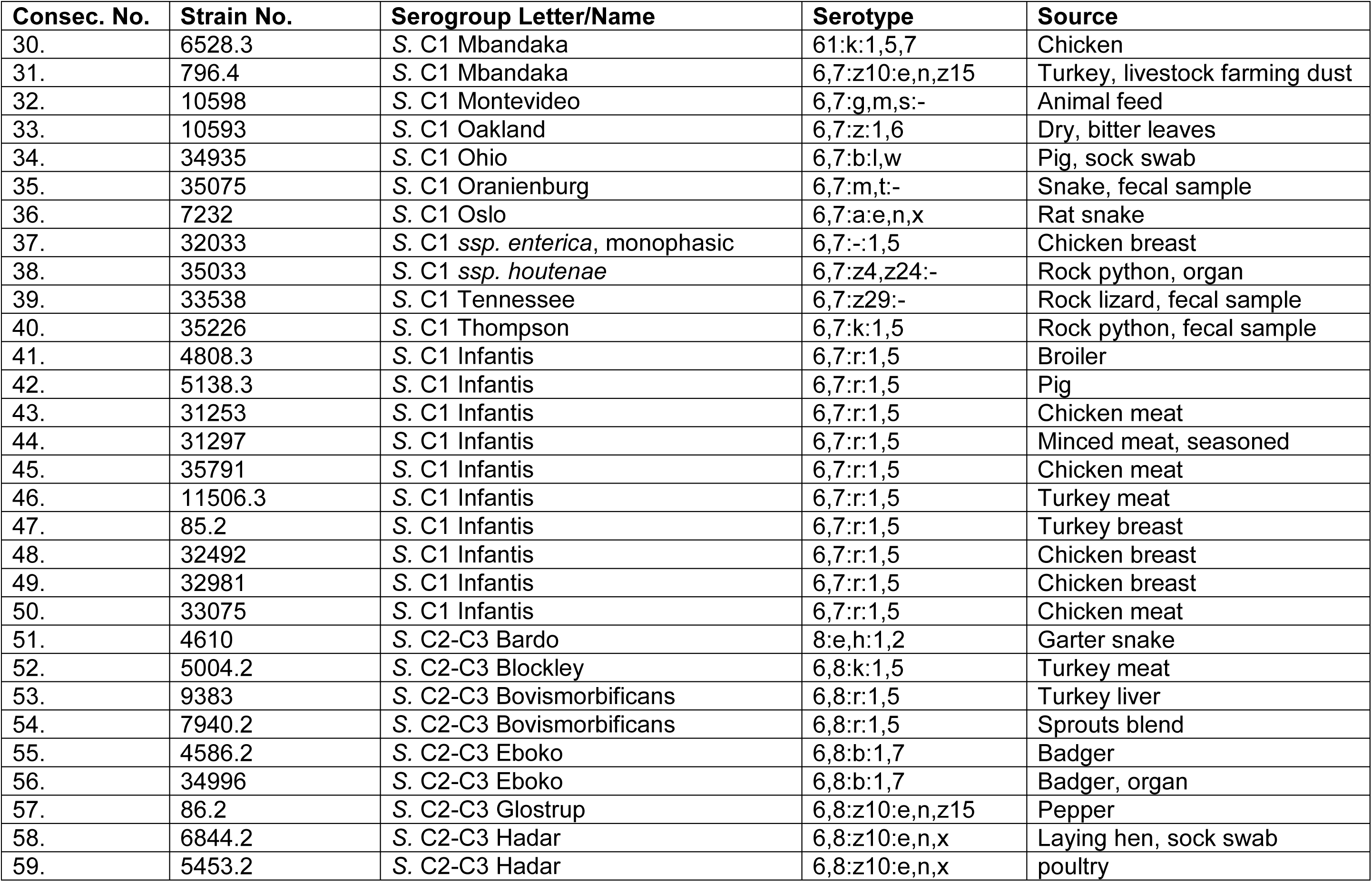
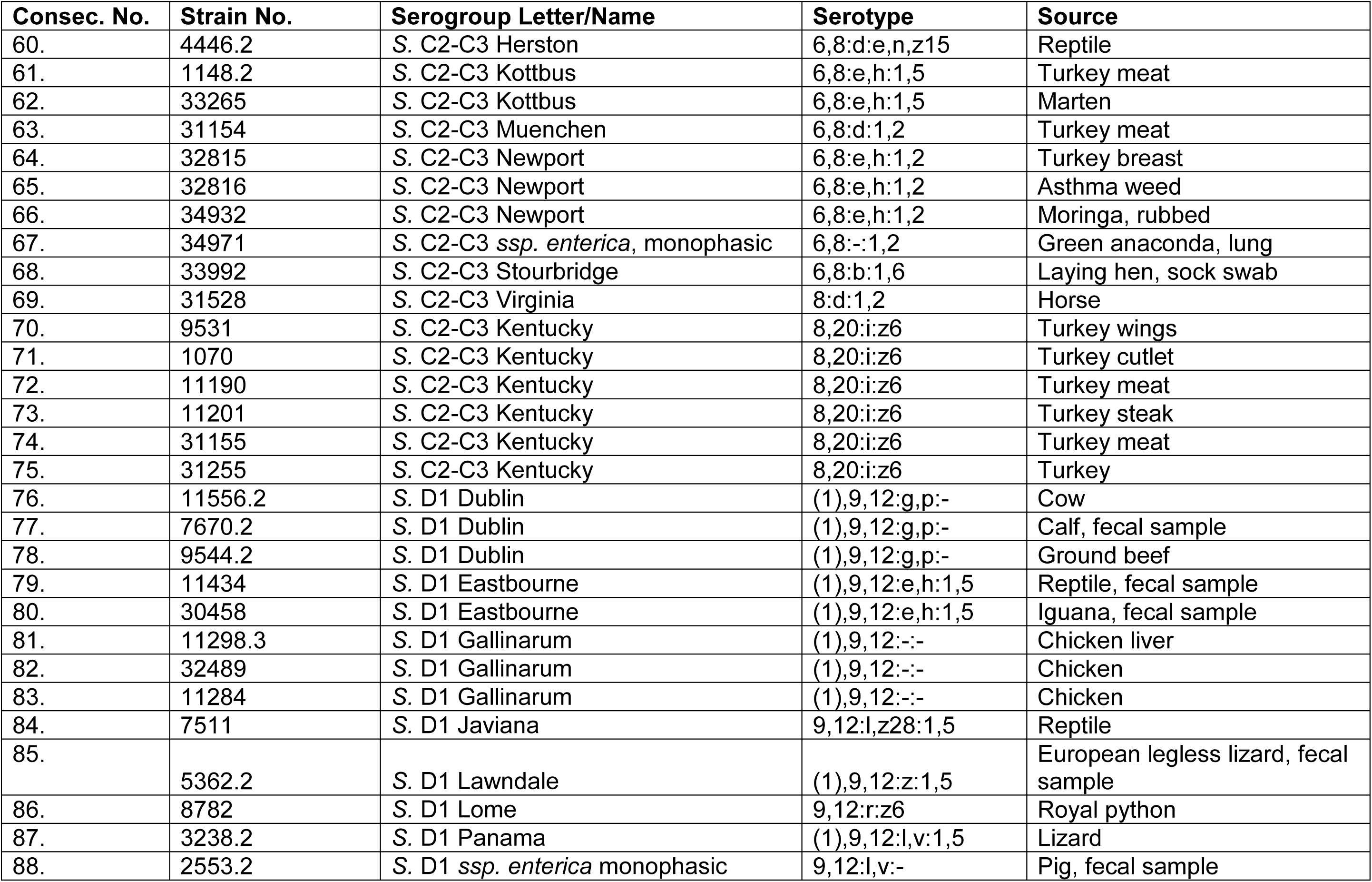
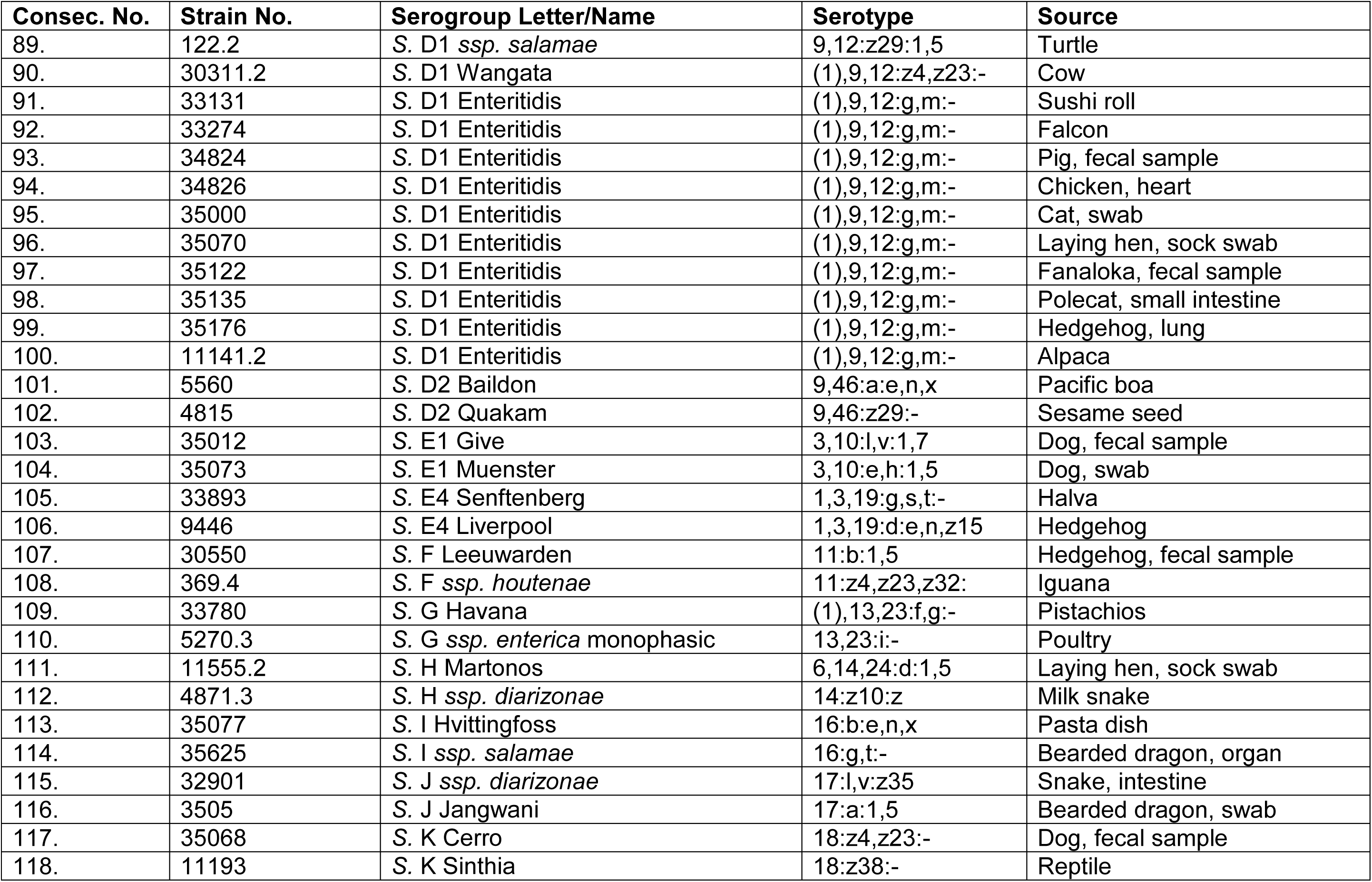
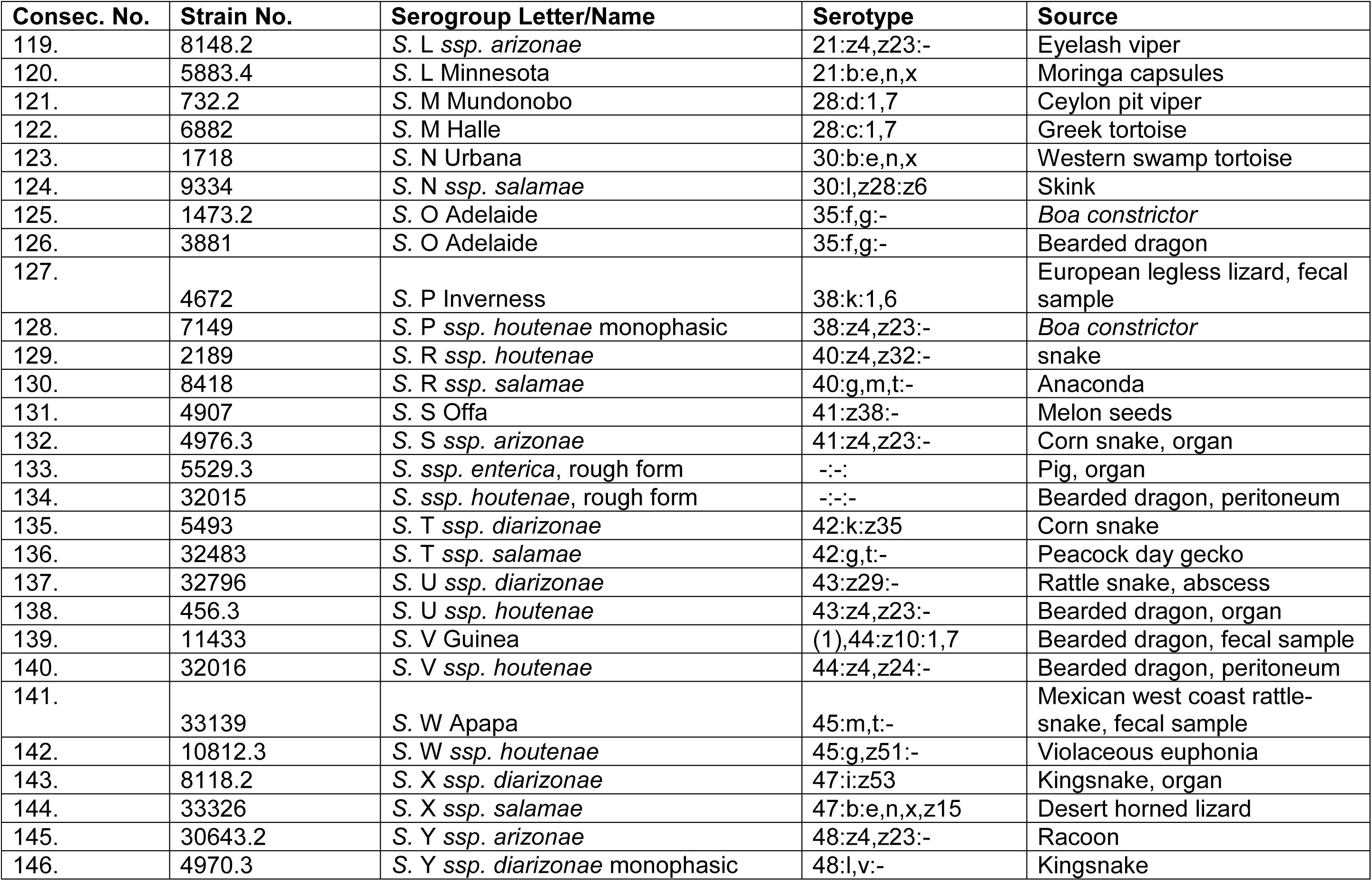
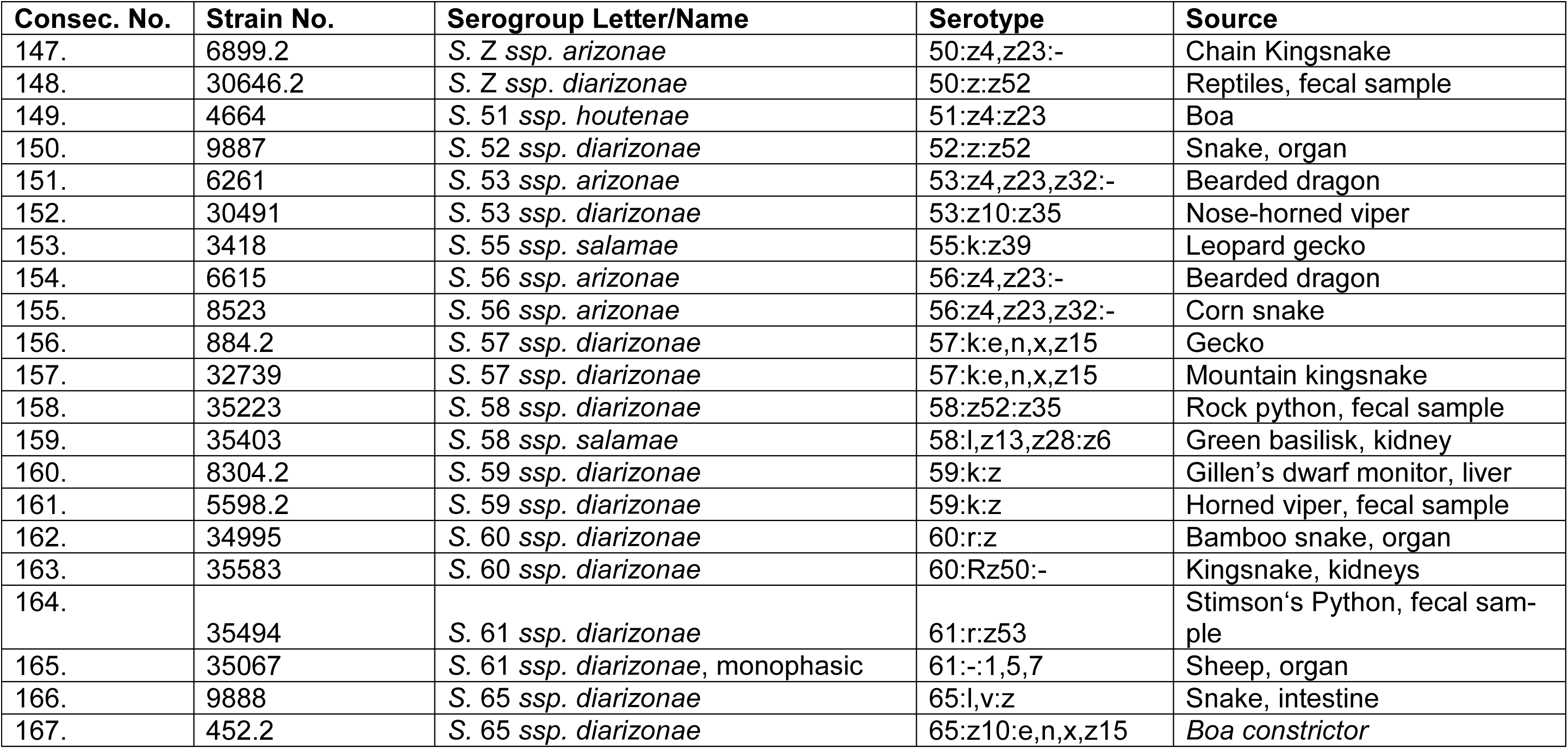
*Salmonella* (*S*.) strains used for the external validation of the Bruker IR-BT^®^ *Salmonella serogroup classifier v3*.

In addition, the *Salmonella serogroup classifier v3* was challenged with a set of *n=*16 salmonella strains from lab proficiency tests performed in our lab from 2006 until 2024 (Tab. 2). This set included *n=*6 strains from serogroup O:4 (B), and *n=*5 strains from each of serogroups O:6,7 (C1) and O:8 (C2-C3).

**Table 2:** Proficiency test *Salmonella* (*S*.) strains used for challenging the Bruker IR-BT^®^ *Salmonella serogroup classifier v3*.

| Consec. No. | Strain No. | Serogroup Letter/Name | Serotype | Year of Proficiency Test |
| --- | --- | --- | --- | --- |
| 1. | **1369** | S. B Bredeney | 4,12:i,v:1,7 | 2006 |
| 2. | **1394** | S. B Typhimurium | (1),4,5,12:i:1,2 | 2006 |
| 3. | **10521** | S. B Typhimurium | (1),4,5,12:i:1,2 | 2015 |
| 4. | **32798** | S. B Typhimurium | (1),4,5,12:i:1,2 | 2020 |
| 5. | **33508** | S. B Typhimurium | (1),4,5,12:i:1,2 | 2021 |
| 6. | **36162** | S. B Typhimurium | (1),4,5,12:i:1,2 | 2024 |
| 7. | **8719.2** | S. C1 Infantis | 6,7:r:1,5 | 2024 |
| 8. | **30307** | S. C1 Infantis | 6,7:r:1,5 | 2017 |
| 9. | **35014** | S. C1 Infantis | 6,7:r:1,5 | 2023 |
| 10. | **2924** | S. C1 Livingstone | 6,7:d:l,w | 2008 |
| 11. | **33269** | S. C1 Oranienburg | 6,7:m,t:- | 2021 |
| 12. | **256.3** | S. C2-C3 Kentucky | 8,20:i:z6 | 2016 |
| 13. | **11723** | S. C2-C3 Litchfield | 6,8:l,v:1,2 | 2017 |
| 14. | **33540** | S. C2-C3 Manchester | 6,8:l,v:1,7 | 2021 |
| 15. | **5298** | S. C2-C3 Manhattan | 6,8:d:1,5 | 2009 |
| 16. | **10696.2** | S. C2-C3 Newport | 6,8:e,h:1,2 | 2016 |

### 2.2 Isolate incubation and IR BT® sample preparation

For the setup of the *Salmonella serogroup classifier v3*, the strains were cultivated on seven different solid media (Columbia agar with 5% sheep blood, Chocolate agar, Tryptose soy agar, Mueller-Hinton agar, XLD agar, Salmonella-Shigella agar and MacConkey agar – Becton Dickinson, Heidelberg, Germany) for 24±2 h at 35±2 °C.

For the validation of the *Salmonella serogroup classifier v3*, the strains (Tab. 1) were incubated in a confluent lawn on Columbia Sheep-Blood (COL-SB) agar (BD Columbia-Agar with 5% Sheep blood by Becton Dickinson, Heidelberg, Germany) for 24±0.5 h at 37 °C. All preparation was performed as outlined previously [29]. Each isolate was cultivated on at least two different COL-SB agar plates and prepared by two different operators (two biological replicates) to simulate preparatory lab variance. Each biological replicate was measured in technical triplicate yielding at least six spectra per isolate.

For the proficiency test strain group challenge, the strains (Tab. 2) were incubated as described above, preparing two technical replicates from two biological replicates each, amounting to four spectra per strain.

IR BT^®^ sample preparation was performed according to the manufacturer’s instructions using the IR Biotyper kit (Bruker Daltonics) [32]. From each agar plate, using a 1 µl loop, cell material was transferred into a 1.5 ml vial containing 50 µl of a 70% (v/v) ethanol solution, and subsequently vortexed to yield a suspension (one suspension replicate per biological replicate). Afterwards, 50 µl deionized water was added to each vial and the suspension mixed by pipetting. Each suspension was transferred onto three spots of a silicon sample plate at 15 µl per spot (three measurement replicates per suspension). After all spots were filled, the sample plate was left to dry within a 37 °C-incubator for approximately 15 min until none of the spots showed visual humidity.

### 2.3 Spectra recording and processing

Absorption spectra were recorded in transmission mode on an IR BT^®^ spectrometer (Bruker Daltonics) between wave numbers 4000 and 500 cm^-1^. The sample plate containing the dried suspension spots was inserted into the instrument’s measurement chamber that is continuously being purged with dried air. The manufacturer’s acquisition method used a resolution of 6 cm^-1^. It performed 32 scans for background and sample spectrum. Zero filling factor was 1. This resulted in 1814 data points in the afore-mentioned wavenumber range from 4000-500 cm^-1^. For post-processing, a “make compatible scalar” function was used interpolating the spectrum, which resulted in a data set with exactly one data point per integer wavenumber, about 3500 points in total.

After measurement, the resulting spectra were subjected to a quality test according to the instructions of the instrument manufacturer [33]. Spectra of poor quality with respect to e.g. minimum and maximum absorbance values and signal-noise-ratio were sorted out so that only spectra with acceptable quality were used for further analyses. Processing of spectra was performed using the IR Biotyper Client software (Bruker Daltonics) [33], applying version V4.0 and using the default settings recommended by the manufacturer described as follows: The spectra were smoothed using the Savitzky-Golay algorithm over 9 data points. Subsequently, the spectra’s second derivative was calculated. Spectra were then cut to the relevant spectral window of 1300-800 cm^-1^, the spectral absorbance region of various oligo- and polysaccharides [34], and hence the suitable spectral range to discriminate bacteria by their serogroups, i.e. outer membrane lipopolysaccharide – based structures, Finally, all spectra were vector-normalized in order to regulate preparation-related variance of biomass and hence absorption.

All qualitatively acceptable spectra were submitted to classification using the *Salmonella serogroup classifier v3* integrated into the IR Biotyper Client software version V4.0. According to the manufacturer, for this machine learning algorithm a linear support vector machine (SVM) predictive model had been built, characterized by using the first 30 Principal Components (PC) and considering the spectral window of 1300-800 cm^-1^ [33], which had been preselected as a wave number range suitable for discriminating between these serogroup classes (Norman Mauder, Bruker, personal communication). For the training of the classifier, *n=*158 *Salmonella* spp. isolates were included, corresponding to *n=*36 O-serogroups.

The serogroup classification result of each spectrum was compared to its expectation due to the serotyping result of the *NRL-Salmonella*. In addition, the resulting number values for both the inlier and outlier score [33], both intrinsic software parameters, were evaluated to assess the classification performance. The inlier and outlier scores indicate whether the recorded spectrum fits into the spectral variance scope of the classifier, thus depicting the reliability of the classification. If the outlier score is lower than the inlier threshold, the spectra are located in the spectral space of the training set. If the outlier score is between inlier and outlier threshold, the spectra are located in the periphery of the spectral space of the training set. A classification score result below the outlier threshold (3.5) indicates a valid classification, while values >3.5 indicate that the sample spectrum is not covered by the spectral variance that was used to establish the classifier, rendering the sample spectrum out of scope and thus questionable.

### 2.4 Spectra exploration method 2D/3D scatter plots

In order to visualize the distribution of the *salmonella* serogroups in the IR spectral space, two- and three-dimensional scatter plot were generated using the IR Biotyper software V4.0 [33]. For this, either the recorded single spectra were used in the case of the proficiency strain set (Tab. 2). For the validation strain set (Tab. 1), in order to decrease visual complexity, all acceptable-quality spectra of one respective isolate were assembled together into one average spectrum. With the respective single or average spectra, 2D and 3D scatter plots were created using the 2^nd^ derivative of the spectra, considering the spectral window of 1300-800 cm^-1^ and the dimensionality reduction algorithm PCA.

### 2.5 Evaluation of results

For quantitative evaluation of results, for each sample spectrum, the following parameters were noted, specified by the IR Biotyper software [33]: 1) the correctness of the serogroup classification in comparison to the actual serotype, and 2) the outlier score. A correct and valid classification was recorded if the attribution matched the actual serotype and the outlier score was ≤3.5. A questionable allocation was noted if the outlier score was >3.5, independent of the actual result assigned. A wrong classification was observed if the allocation did not match the actual serotype given that the outlier score was ≤3.5, respectively (supplementary material S1, S2, S3, S4).

The validation concept performed was inspired by the BVL Guidelines for Validating Species Identifications using MALDI-ToF-MS in a single laboratory [27, 28]. The evaluation of the external validation for the *Salmonella* serogrouping method using the IR Biotyper was performed as a so-called *targeted identification.* In order to assess the inclusivity, at least 20 strains of the target parameter needed to be validated with a true positive rate of ≥95% and a false negative rate of ≤1%. At the same time, the exclusivity was assessed by at least 30 validated strains of the non-target parameters with a true negative rate of ≥99% and a false positive rate of ≤1%. In this case, the target parameter was the respective serogroup under consideration, whereas the entirety of the other serogroups together with the Noah’s ark strain group represented the non-target parameter. Thus, this evaluation was carried out for each desired target parameter separately within the study set under consideration (Fig. 1-4). As an additional requirement, at least 90% of both the target and the non-target parameter spectra had to be classified, i.e. a maximum of 10% of the spectra was admitted to result as questionable.

**Fig 1:**
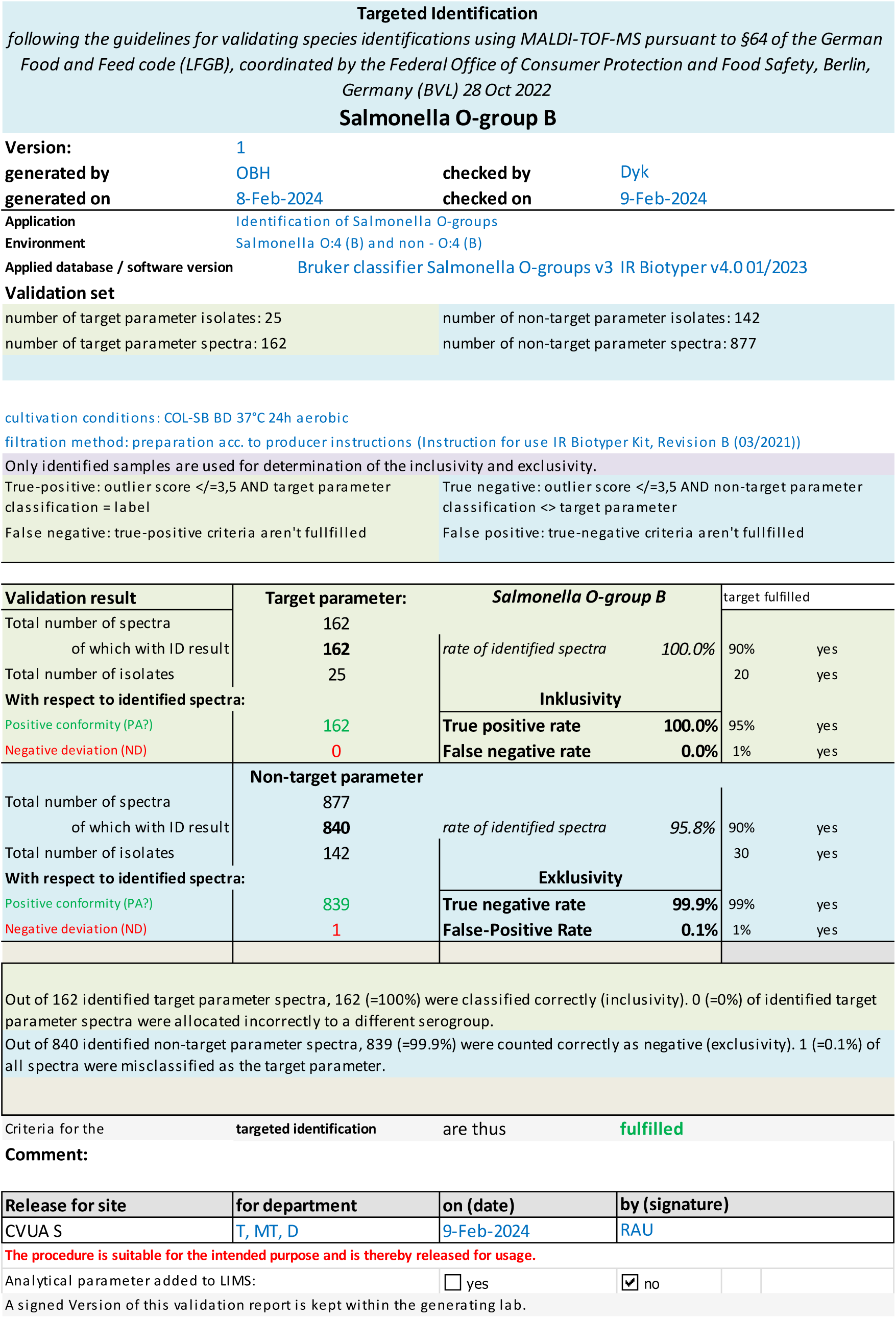
Summary report of the targeted identification of *Salmonella* serogroup **B (O:4)** following the guidelines for validating species identifications using MALDI-TOF-MS pursuant to §64 of the German Food and Feed code (LFGB), coordinated by the Federal Office of Consumer Protection and Food Safety, Berlin, Germany (BVL) 28 Oct 2022.

## 3 Results and Discussion

### 3.1 Classifier set-up

To build the *Salmonella serogroup classifier v3*, a support vector machine (SVM) predictive model was been built by means of the IR Biotyper software 4.0. For the training of the classifier, *n=*158 *Salmonella* spp. isolates were included, corresponding to 36 O-serogroups and cultivated on seven different, widely used culture media. For the verification of the classifier, different internal (belonging to the study design) and external (independent) datasets were used [24].

### 3.2 Classifier validation

The *Salmonella serogroup classifier v3* was subsequently validated by a procedure inspired by the Guidelines for Validating Species Identifications by MALDI ToF MS [28], as previously successfully applied for the validation of a *Listeria monocytogenes* serogroup classifier [29]. For this, a total of *n=*630 spectra from *n=*100 strains from four different target groups (Tab. 1) were classified using the above-mentioned classifier and all results noted (suppl mat S1, S2, S3, S4). The respective outgroup to each target parameter consisted of both the other three respective serogroups under consideration and a large group of *n=*409 spectra from a large set of *n=*67 strains from *n=*35 different non-targeted serogroups in addition. Since two different serotype representatives were randomly chosen per serogroup, this group was termed “Noah’s Ark strains”. The Noah’s Ark strains served as a background to challenge the *Salmonella serogroup classifier v3* with a large set of diverse outgroup strains that might occur rarely, but nevertheless realistically in daily lab life.

Concerning the target parameter *serogroup O:4 (B)* (Fig. 1), all *n=*162 spectra were correctly identified as O:4 indeed. Out of *n=*877 non-target parameter spectra, *n=*840 (=95.8%) were classified, all but one spectrum being correctly identified as not belonging to serogroup O:4. Therefore, this parameter’s sensitivity was 100% and exclusivity was 99.9%.

Similar results were noted for both serogroups O:6,7 (C1) and O:9 (D1): With respect to *serogroup O:6,7 (C1)* (Fig. 2), *n=*148 out of *n=*150 target parameter spectra were correctly allocated to serogroup C1, corresponding to a true-positive rate of 99.3%. 96.1% of the non-target parameter spectra were identified and their classifications correct in each case, amounting to a true-negative rate of 100%.

**Fig 2:**
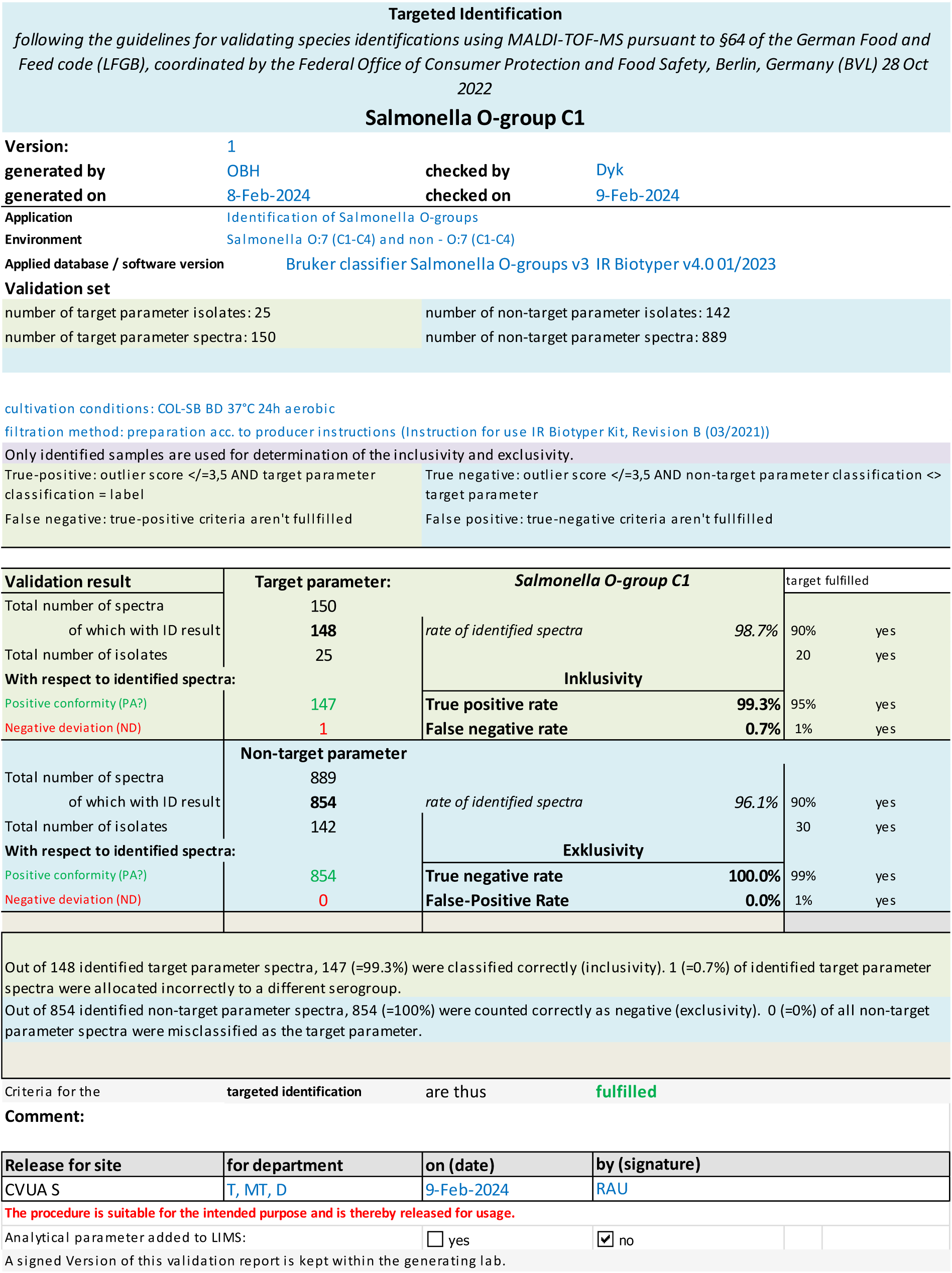
Summary report of the targeted identification of *Salmonella* serogroup **C1 (O:7)** following the guidelines for validating species identifications using MALDI-TOF-MS pursuant to §64 of the German Food and Feed code (LFGB), coordinated by the Federal Office of Consumer Protection and Food Safety, Berlin, Germany (BVL) 28 Oct 2022.

Regarding *serogroup O:9 (D1)* (Fig. 3), both the rate of identified target parameter spectra and their true-positive rate were 100% each. With respect to the non-target parameter, the exclusivity was 99.3%, i.e. from *n=*843 identified spectra, all *n=*6 spectra from isolate CVUAS 30643.2 (*S. enterica ssp*. *arizonae* 48:z4,z23:-) were incorrectly assigned to the target parameter. Nevertheless, as the resulting false-positive rate of 0.7% is below the threshold of 1%, this target parameter *serogroup O.9 (D1)* was considered as fully validated.

**Fig 3:**
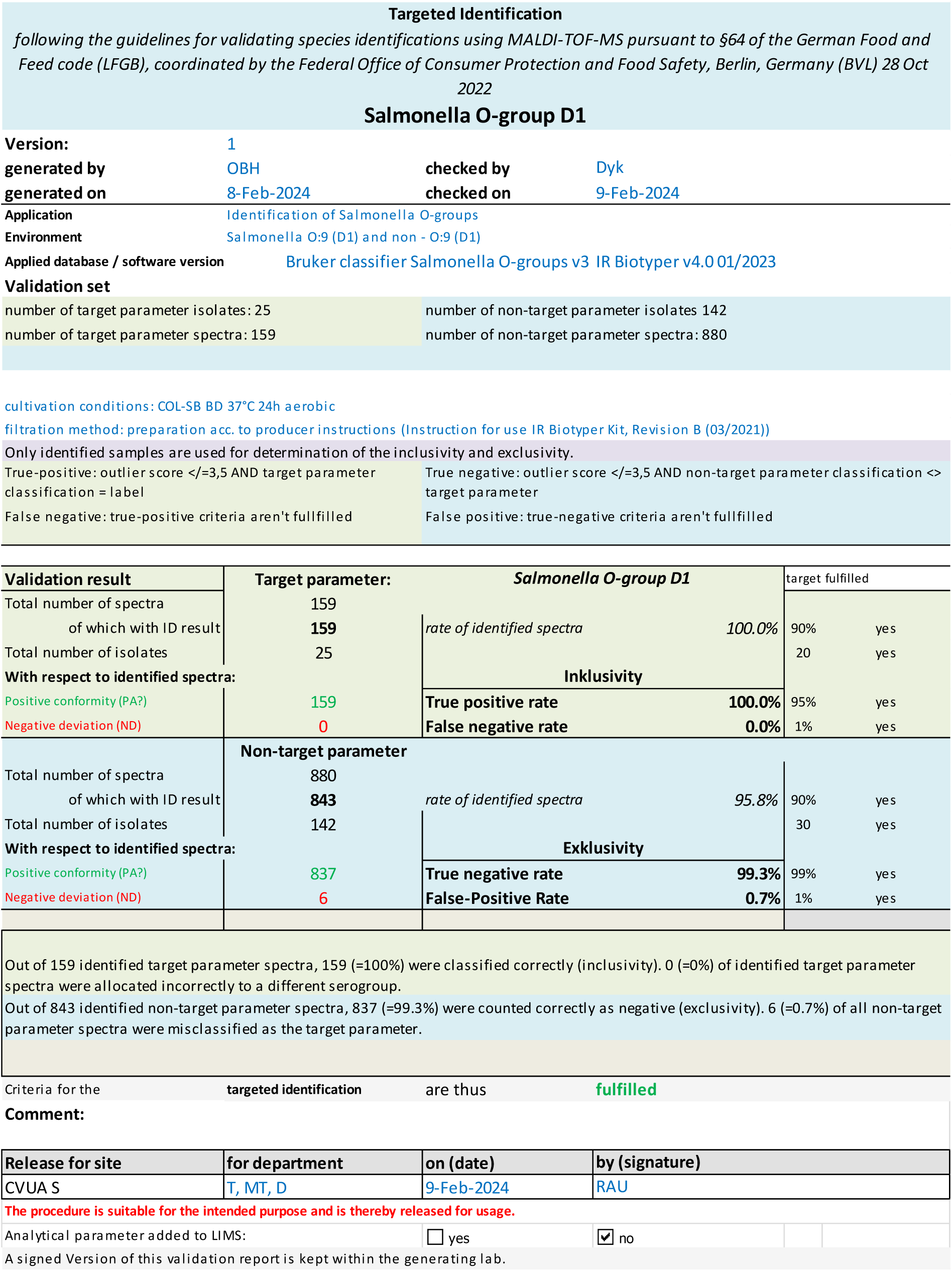
Summary report of the targeted identification of *Salmonella* serogroup **D1 (O:9)** following the guidelines for validating species identifications using MALDI-TOF-MS pursuant to §64 of the German Food and Feed code (LFGB), coordinated by the Federal Office of Consumer Protection and Food Safety, Berlin, Germany (BVL) 28 Oct 2022.

In this validation, the results of the target parameters *serogroup O:4, serogroup O:6,7* and *serogroup O:9* were fully in line with the adapted requirements of the national guidelines [28] subsidiarily used for validation and can therefore be considered as successfully validated by this challenging scheme (Figs. 1-3).

Respecting the target parameter *serogroup O:8 (C2-C3)* (Fig. 4), while 96.1% of the target parameter’s spectra were correctly assigned, the classifier misallocated all *n=*6 spectra from strain CVUAS 4586 (*S.* Eboko 6,8:b:1,7) to different serogroups. This incorrect classification resulted in a false-negative rate of 3.9% and thus, this parameter did not meet the requirements of the adapted Guidelines for Validating Species Identifications [28]. Regarding the non-target parameter spectra, all of the 96.4% identified spectra were correctly classified as not belonging to serogroup O:8.

**Fig 4:**
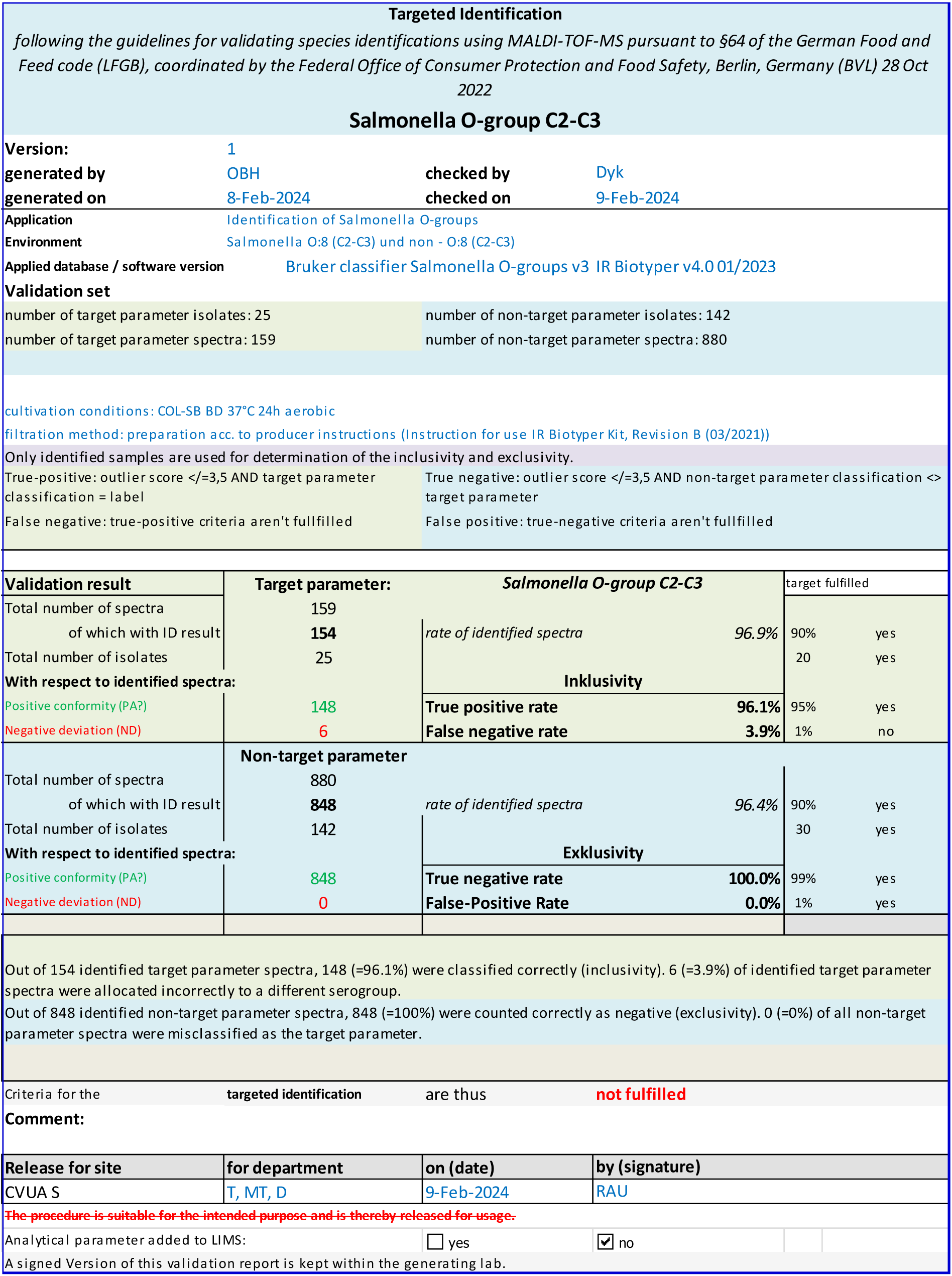
Summary report of the targeted identification of *Salmonella* serogroup **C2-C3 (O:8)** following the guidelines for validating species identifications using MALDI-TOF-MS pursuant to §64 of the German Food and Feed code (LFGB), coordinated by the Federal Office of Consumer Protection and Food Safety, Berlin, Germany (BVL) 28 Oct 2022.

The false-negative identification of strain CVUAS 4586 suggests that the classifier in its current form does not cover the full biodiversity of this particular serogroup O:8. Therefore, the inclusion of more strains into the training set would most likely improve the classifier’s performance and render it more powerful to faithfully recognize this particular serogroup.

The quality of differentiation performance found in the current study has been observed before: Results for true positive rates were in a range previously described for *Salmonella* serovar differentiation by FTIR spectroscopy [15, 18, 20–22, 24, 25]. Recently, Napoleoni *et al*. noted a sensitivity of 100% for serogroup B, 91.5% for serogroup C1 and 98.2% for serogroup D1 [26]. With respect to the proficiency test strain set (Tab. 2) used to challenge the classifier, each of the spectra was correctly allocated to its respective serogroup.

### 3.3 Visualization of spectra in the spectral space

In the 2D scatter plot depicting all *n=*64 proficiency strain set single spectra (Tab. 2) from this study with respect to their most discriminatory Principal Component values PC 1 and 2, the separation between the different classes is clearly visible (Fig. 5) and demonstrates the differentiation capability of the trained classifier for strains belonging to the specific serogroups under investigation.

**Fig 5:**
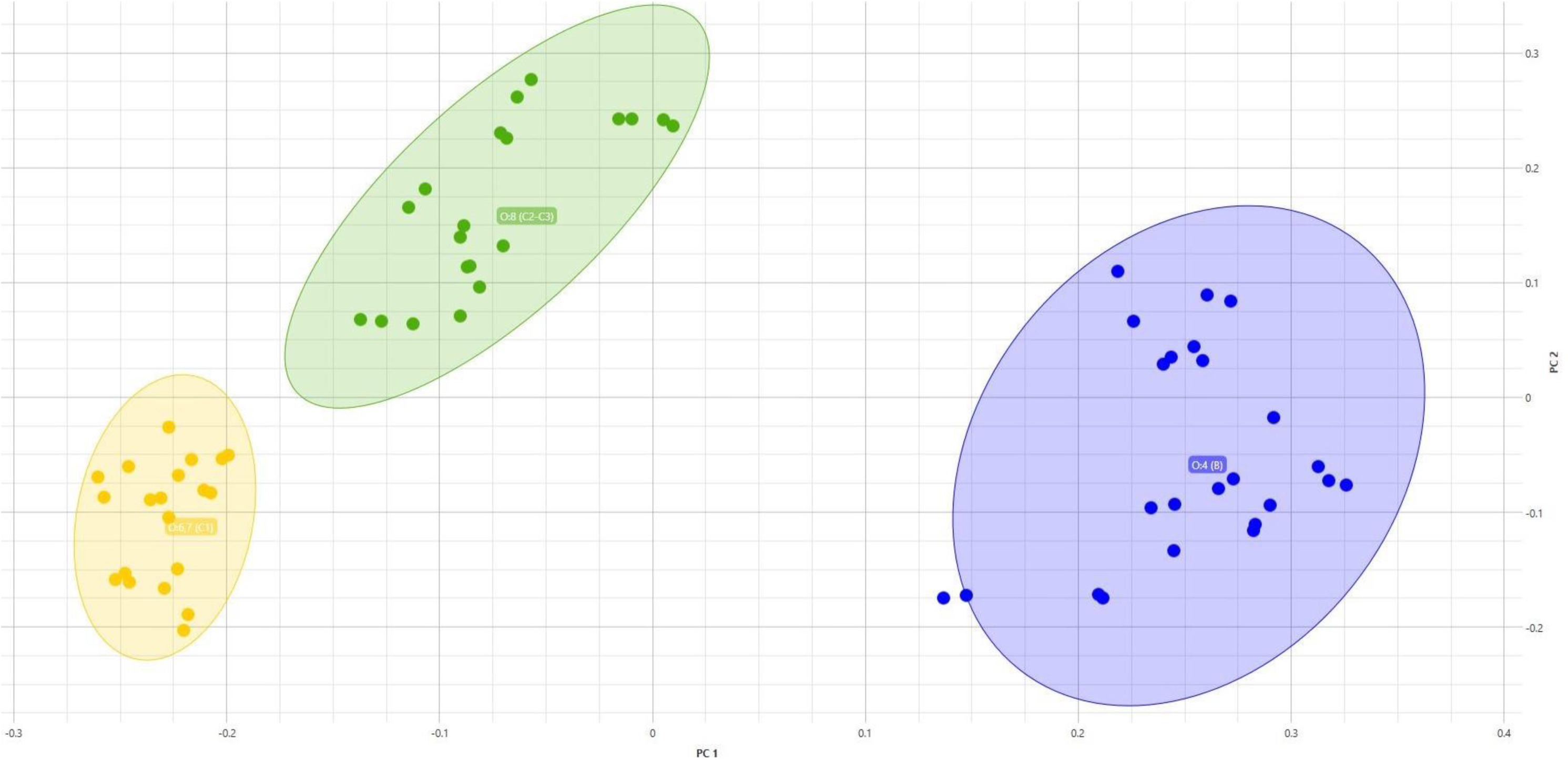
2D-scatter plot of the proficiency test strains’ single spectra (Tab. 2), 2^nd^ derivative, spectral range 1300-800 cm^-1^, dimensionality reduction algorithm PCA, displaying Principal Components PC 1 and PC 2.

The 3D scatter plot showing all *n=*167 validation strain set average spectra (Tab. 1) in relation to their most prominent Principal Components PC1, PC2 and PC3 shows a more intricate picture (Fig. 6): While the spectra clusters of serogroups O:9 and O:4 are comparatively large and well separated from the other spectra within the spectral space spread out by PC 1 to PC 3, the spectra clusters of serogroups O:6,7 and O:8 are much more confined and thus depict less spectral variance. The spectra from serogroups O:6,7 and O:8 also intermingle more with the Noah’s Ark strain spectra in this visualization. Nevertheless, as the external validation demonstrates, the AI-based classifier was trained successfully in working out the discriminatory features of serogroup O:6,7 and with one deviating strain also of serogroup O:8.

**Fig 6:**
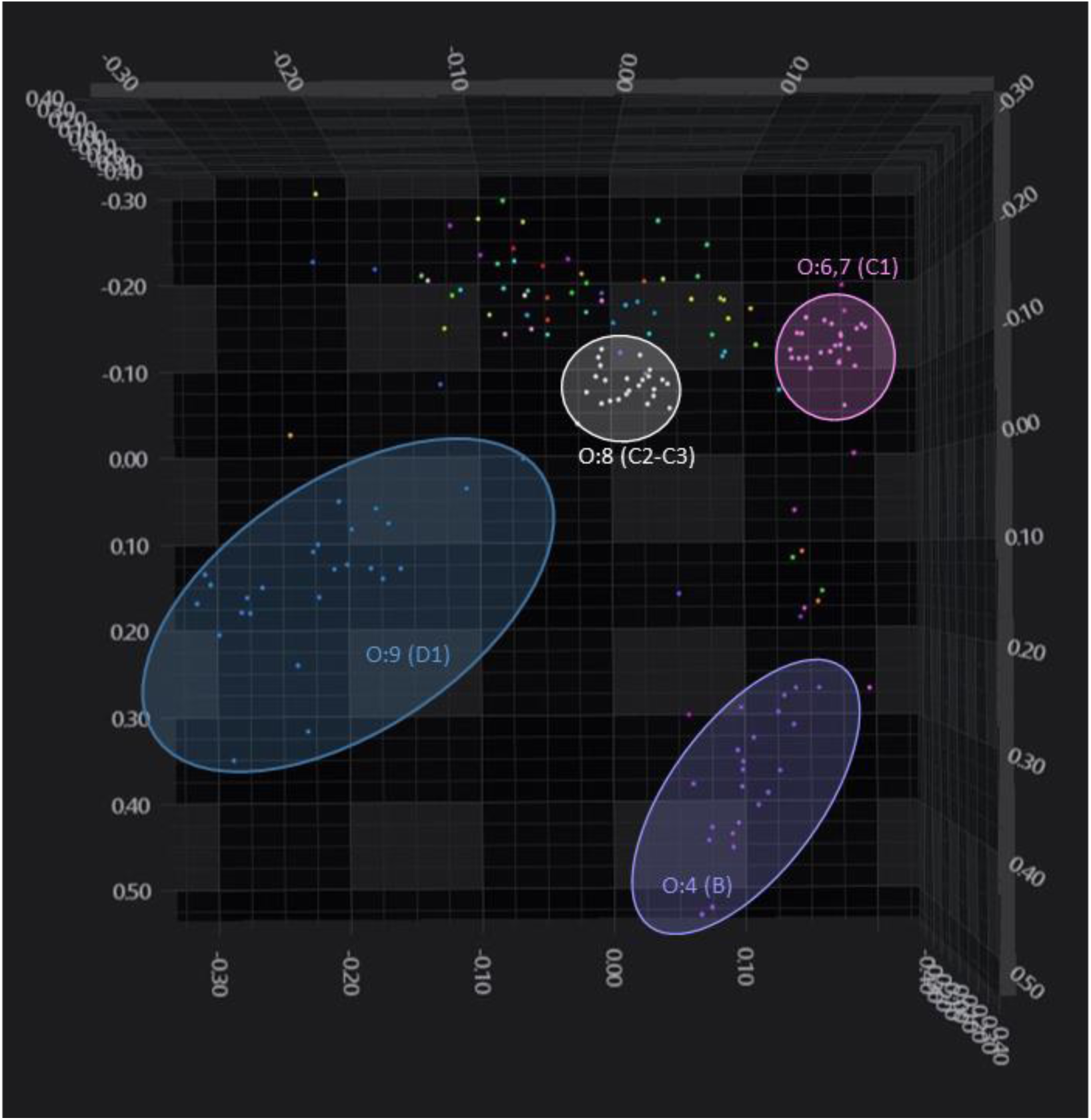
3D-scatter plot of the validation strains’ average spectra (Tab. 1), 2^nd^ derivative, spectral range 1300-800 cm^-1^, dimensionality reduction algorithm PCA, displaying Principal Components PC 1, PC 2 and PC 3. Colorful non-marked dots denote Noah’s ark strains.

## 4 Concluding Remarks

The suitability of the *Salmonella serogroup classifier v3* to distinguish between different *Salmonella* serogroups has thus been demonstrated for serogroups O:4, O:6,7 and O:9, three very frequently occurring serogroups of our daily lab practice. The fourth serogroup under investigation, O:8, failed meeting the adapted Guidelines [28] by only one strain out of *n=*25 target parameter strains, suggesting an improvement being well possible. The classifier can now be used within our accredited lab work-flow.

The transfer of the adapted guidelines for the validation of species identifications by MALDI ToF MS [28] provides users of any spectroscopy or spectrometric classification method transparent guidance to carry out any external validations, thereby simplifying the implementation of new methods.

The Guidelines demand large datasets be analyzed with respect to both the target parameter and the non-target parameter, thereby emulating conditions close to reality in an analytical laboratory with a high sample throughput. The classification method under investigation was thus challenged and its performance evaluated under demanding conditions in order to obtain a robust and reliable result. Any classifier passing this performance test has therefore been proven to achieve the desired goal. Recently, a *Listeria* serogroup classifier has been proven by this method to reliably distinguish between different serogroups of *Listeria monocytogenes* [29].

As a future perspective, a serovar specific sub-classifier would enable the discrimination among selected serovars of the same serogroup. For instance, the specific determination of the very frequent Serovars *S.* Typhimurium (O:4), *S.* Infantis (O:6,7) and *S.* Enteritidis (O:9) would be both helpful to save two days’ time for analysis and decrease the amount of rabbit sera, both necessary for conventional serotyping. A faster analytical approach by FTIR spectroscopy over conventional serotyping would be an attractive method, especially for livestock analysis with frequently repetitive isolates, such as e.g. salmonella vaccination strains in chicken barns.

## Acknowledgements

The authors wish to thank P. Bayan and L. Loeffler, both CVUA Stuttgart, for their excellent technical assistance.

## Statements

MC is a Bruker Daltonics employee. The other authors declare that they have no known competing financial interests or personal relationships that could have appeared to influence the work reported in this paper. While Bruker Daltonics, Germany, set up the commercial classifier *Salmonella serogroup classifier v3*, Bruker Daltonics had no influence on the design of the validation nor in the collection, analyses or interpretation of the validation data. This study does not involve any human participants or animal experiments.

## Data availability

Data will be made available on request. Metadata of the isolates used in the validation are listed in the MALDI-UP catalogue, https://maldi-up.ua-bwl.de, including functional spreadsheet data files of all validations performed in the context of this study.

## Supplementary material

Supplementary data associated with this article can be found at the end of this manuscript.

